# Characterization of transcriptional heterogeneity and novel therapeutic targets using single cell RNA-sequencing of primary and circulating Ewing sarcoma cells

**DOI:** 10.1101/2024.01.18.576251

**Authors:** Andrew Goodspeed, Avery Bodlak, Alexis B. Duffy, Sarah Nelson-Taylor, Naoki Oike, Timothy Porfilio, Ryota Shirai, Deandra Walker, Amy Treece, Jennifer Black, Nathan Donaldson, Carrye Cost, Tim Garrington, Brian Greffe, Sandra Luna-Fineman, Jenna Demedis, Jessica Lake, Etienne Danis, Michael Verneris, Daniel L Adams, Masanori Hayashi

## Abstract

Ewing sarcoma is the second most common bone cancer in children, accounting for 2% of pediatric cancer diagnoses. Patients who present with metastatic disease at the time of diagnosis have a dismal prognosis, compared to the >70% 5-year survival of those with localized disease. Here, we utilized single cell RNA-sequencing to characterize the transcriptional landscape of primary Ewing sarcoma tumors and surrounding tumor microenvironment (TME). Copy-number analysis identified subclonal evolution within patients prior to treatment. Primary tumor samples demonstrate a heterogenous transcriptional landscape with several conserved gene expression programs, including those composed of genes related to proliferation and EWS targets. Single cell RNA-sequencing and immunofluorescence of circulating tumor cells at the time of diagnosis identified TSPAN8 as a novel therapeutic target.

## Introduction

While the intensification of conventional multi-modal therapy has significantly improved the survival rate for localized Ewing sarcoma from less than 10% in the surgery-only era to approximately 70%, many patients still die from metastatic disease^1, 2, 3^. With intensive multi-modal therapy using chemotherapy, radiation, and surgery, many patients, even those with initial metastatic disease achieve a radiographic remission; however, metastatic relapse after a remission frequently occurs. Accordingly, an initial good response to conventional therapy does not guarantee a cure.

While the Ewing sarcoma genomic landscape involves few mutations, the pathognomonic t(11;22) EWS-FLI1 translocation drives 90% of the disease^4, 5, 6, 7^. By studying EWS-FLI1 transcriptional regulation, over 1,000 genes have been identified to be directly or indirectly induced or repressed^8, 9, 10, 11^. While these results have spurred a significant effort focused on EWS-FLI1-induced genes that might be targeted to inhibit tumor progression, clinical translation has been difficult^12, 13^. While EWS-FLI1 expression is inarguably essential to the survival of Ewing sarcoma cells, emerging experimental and clinical evidence suggests that Ewing sarcoma tumors have significant intra-tumoral EWS-FLI1 transcriptional heterogeneity^14, 14, 15^. Additionally, conditional knockdown systems of EWS-FLI1 have indicated that low-level EWS-FLI1 promotes a pro-metastasis cell phenotype, compared to high expression^14^. A major unmet need is to better understand tumor heterogeneity and how it potentially contributes to poor prognosis, with the goal of identifying therapeutic targets that are essential to metastasis and recurrence.

The understanding of the Ewing sarcoma tumor microenvironment (TME) is still evolving. Previous work in this cancer type has suggested the TME is infiltrated by immunosuppressive myeloid cells, functionally impaired antigen presenting cells, especially conventional dendritic cells, and T cells with features suggestive of dysfunction^16, 17^. Ewing sarcoma tumor cells have been shown to lack expression of HLA class I molecules, potentially leading to decreased T cell recognition ^16, 18^. Furthermore, T cells in Ewing sarcoma cells have been indicated to have high LGALS3 and LAG3 expression, further suggesting dysfunction^16, 17^. Tumor-TME interaction has been suggested to involve Wnt signaling^19^, USP6^20^, and VEGF^21, 22^ but whether these interactions are present beyond cell line and mouse PDX models is unclear.

Despite these advances in the investigation of Ewing sarcoma tumor biology, clinical support and translation has been poor. For instance, the knowledge of intra-tumoral transcriptional activity in untreated patient tumors is limited, and we know very little about how this may drive treatment resistance or clonal evolution. Here, we used single cell RNA-sequencing (scRNA-seq) to investigate the transcriptional heterogeneity of untreated Ewing sarcoma patient tumors, as well as circulating tumor cells (CTCs). We found that patient tumors contain heterogenous clusters of tumor cells with distinct genetic expression programs, which correlate with patient outcomes. Evaluation of primary tumors, CTCs, and the TME revealed several potential therapeutic targets.

## Results

### Single cell RNA-seq reveals transcriptional heterogeneity in Ewing Sarcoma primary tumors

To characterize the intratumoral landscape of Ewing sarcoma, we performed droplet-based scRNA-seq on biopsies of primary tumor samples from seven patients with newly diagnosed Ewing sarcoma (Supplementary Fig. 1A). Six of the tumors had an EWS-FLI1 translocation with the other containing EWS-ERG (Table 1). Additionally, 3 tumors had mutations in previously associated Ewing sarcoma-altered genes^5, 6^, including 1 with the most frequently mutated gene, STAG2, demonstrating clinical relevance of these samples (Supplementary Fig. 1B). All samples were from bone primary tumors and were obtained prior to any systemic therapy. One patient presented with metastasis, and the others with localized disease. After doublet removal and filtering of low-quality cells, the scRNA-seq dataset consisted of 6228 cells with 98936678 Unique Molecular Identifiers (UMIs) across 22518 genes (Supplementary Fig. 1C-D, Supplementary Table 1).

**Table 1.**

| ID | Patient Data | Location | Metastases | Fusion | Tumor Karyotype | Recurrence | OS (months) | Samples |  |
| --- | --- | --- | --- | --- | --- | --- | --- | --- | --- |
|  |  |  |  |  |  |  |  | Tumor | CTC Blood |
| 001 | 13y M | L fibula | - | EWS-ERG | EWS-ERG only | - |  | + | + |
| 002 | 19y F | Pleural mass | + (spine) | EWS-FLI1 |  | + (local progression on therapy) | 12 | - | + |
| 005 | 15y M | R iliac crest | - | EWS-FLI1 | Hyperploidy: 5, 8, 1q | + (local, 9 months off therapy) |  | + | + |
| 006 | 8y M | Paraspinal | - | EWS-FLI1 |  | - |  | - | + |
| 009 | 15y F | Sacrum | + (R humerus) | EWS-FLI1 | Hyperploidy: 2, 4, 12, 16, 20, 21 | + (local, 7 months off therapy) | 29 | - | + |
| 010 | 15y M | R humerus | - | EWS-FLI1 | EWS-FLI1 only | - |  | + | + |
| 012 | 17y F | Cutaneous (R thigh) | - | EWS-FLI1 |  | - |  | - | + |
| 038 | 9y M | L ileum | - | EWS-FLI1 | Hyperploidy: 8 | + (metastatic, 3 months off therapy, lung) | 24 | + | + |
| 051 | 19y M | R ulna | + (liver, spine, sacrum, ilium, femurs) | EWS-FLI1 | Hyperploidy: 2, 8, 13, 21 | + (metastatic 2 months off therapy, extensive disease in bones and brain) | 16 | + | - |
| 061 | 6y M | R scapula | - | EWS-FLI1 | Hyperploidy: Clone 1: 2, 8, 20 clone 2: Additional gain of 5, 12, 13 | - |  | + | - |
| 066 | 14y M | L scapula | - | EWS-FLI1 | EWS-FLI1 only | - |  | + | - |

Single cells were visualized using uniform manifold approximation and projection (UMAP) followed by unbiased clustering (Supplementary Fig. 2A). We classified nine major cell types (Fig. 1A-B, Supplementary Fig. 2B, Supplementary Table 2). In addition to Ewing sarcoma, we found the tumor microenvironment (TME) to consist of myeloid cells, endothelial cells, T cells, B cells, osteoclasts, and matrix and inflammatory cancer associated fibroblasts (CAFs) (Fig. 1C-D). Many of these cell types were present in all tumor samples, although inflammatory CAFs were only observed in patient 001 (Supplementary Fig. 2C).

**Figure 1.**
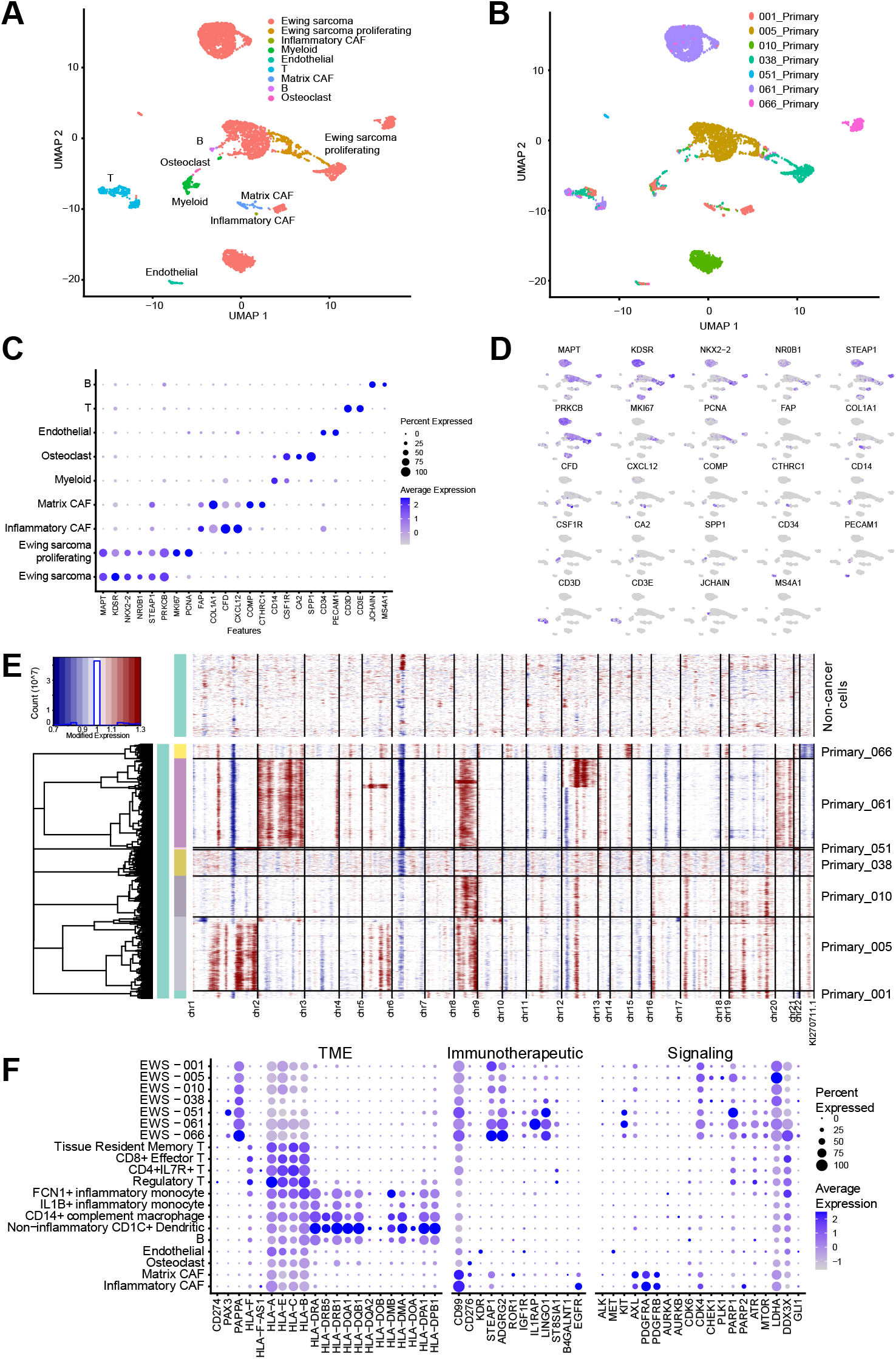
scRNA-seq of Ewing sarcoma primary tumors reveals several cell types including cancer cells. UMAP projections labeled by identified cell type (A) or the sample source (B). Expression of canonical marker genes across cell types are shown as a dotplot (C) or UMAP (D). Inferred copy-number alteration estimated in the identified cancer cells separated by sample with all other cell types serving as control (E). Expression of select genes within Ewing sarcoma tumor cells from specific patients and aggregate cell types (F).

Ewing sarcoma cells were identified through several means. These cells expressed high levels of canonical Ewing sarcoma-associated genes such as such as MAPT^23^, KDSR^24^, NKX2-2^25, 26^, NR0B1^33, 27, 28^, STEAP1^29, 30^, and PRKCB^31^ (Fig 1C-D, Supplementary Fig. 2D). The classification of these cells as malignant was further supported by single cell variational aneuploidy analysis (SCEVAN) (Supplementary Fig. 2E) and by the enrichment of several EWS-FLI1 signature gene sets (Supplementary Fig. 2F). Among the Ewing sarcoma cells, a proliferating population was identified by the expression of MKI67 and PCNA (Fig. 1C-D).

A benefit of scRNA-seq is the ability to characterize several forms of tumor heterogeneity. Here, we utilized inferCNV to infer genome-wide copy number alterations (CNAs) across the seven primary tumors^32^ (Fig. 1E, Supplementary Fig. 2G). Previous Ewing sarcoma studies have identified several common CNAs, including gains in chr 1q, chr 2, chr 5, chr 7, chr 8, chr 12, chr 20, and loss of chr 9p and chr 16q^6, 33, 34, 35, 36, 37, 38^. We identified several of these alterations such as gains of chr1q, chr2, chr5, chr8, chr12, and chr20. These findings closely corroborated with the clinical karyotyping of these primary tumors (Table 1).

The expression of previously reported preclinical and clinical therapeutic targets for Ewing sarcoma in our data set (Fig. 1F). TME-, cell surface-, and cell intrinsic signaling-related targets were plotted across cell type and within tumor cells by sample. Genes comprising the surfaceome of Ewing sarcoma were also plotted (Supplementary Fig. 2H)^39^. We found that compared to other cell types in the primary tumor, Ewing cells across all samples express moderate to high levels of PAPPA but low levels of most HLA genes. For immunotherapeutic targets, recent efforts have identified several candidates such as IL1RAP, CD99, STEAP1, ADGRG2, ENPP1, CDH11, and LINGO1^39, 40, 41^. We show that several proposed immunotherapeutic targets such as ADGRG2, IL1RAP, and LINGO1 are uniquely and highly expressed in Ewing sarcoma (Fig. 1F, Supplementary Fig. 2H).. CD99 and STEAP1 are also highly expressed in Ewing sarcoma cells, although there is significant expression in CAFs as well. While mRNA expression and cell surface expression does not always correlate^42^, it is quite notable that the expression of many of these genes is heterogenous across tumor samples, especially for IL1RAP and LINGO1. We found that many of the potential cell signaling-related therapeutic targets are expressed heterogeneously within the tumors, and across patient samples, including CDK4, PARP1, and DDX3X. RNA alone may not correlated with activity but our data confirms what has been widely observed in the clinic; despite the genetic makeup that suggests otherwise, Ewing sarcoma does not seem to have a single dependency that can be targeted with a single agent.

### Ewing sarcoma tumors demonstrate transcriptional heterogeneity

Next, we sought to define heterogeneity of transcriptional programs Ewing sarcoma tumors. The tumor cells identified in Fig. 1 were isolated, integrated across patients, and then we performed non-negative matrix factorization (NMF) analysis in a sample-specific manner to identify intratumor gene expression programs (GEPs). This revealed a total of 25 GEPs (Supplementary Table 3), which were grouped into six main meta-programs through hierarchical clustering (Fig. 2A, Supplementary Table 4). Most of the expression programs were represented in multiple tumors, suggesting that they represent diverse functions shared among patients. Based on the gene expression and gene set over-representation analysis within each merged GEP, we termed the 6 merged GEPs MAPK, EWS, Proliferation, UPR (unfolded protein response), Ribosome, and OxPhos (oxidative phosphorylation) (Fig. 2B-E).

**Figure 2.**
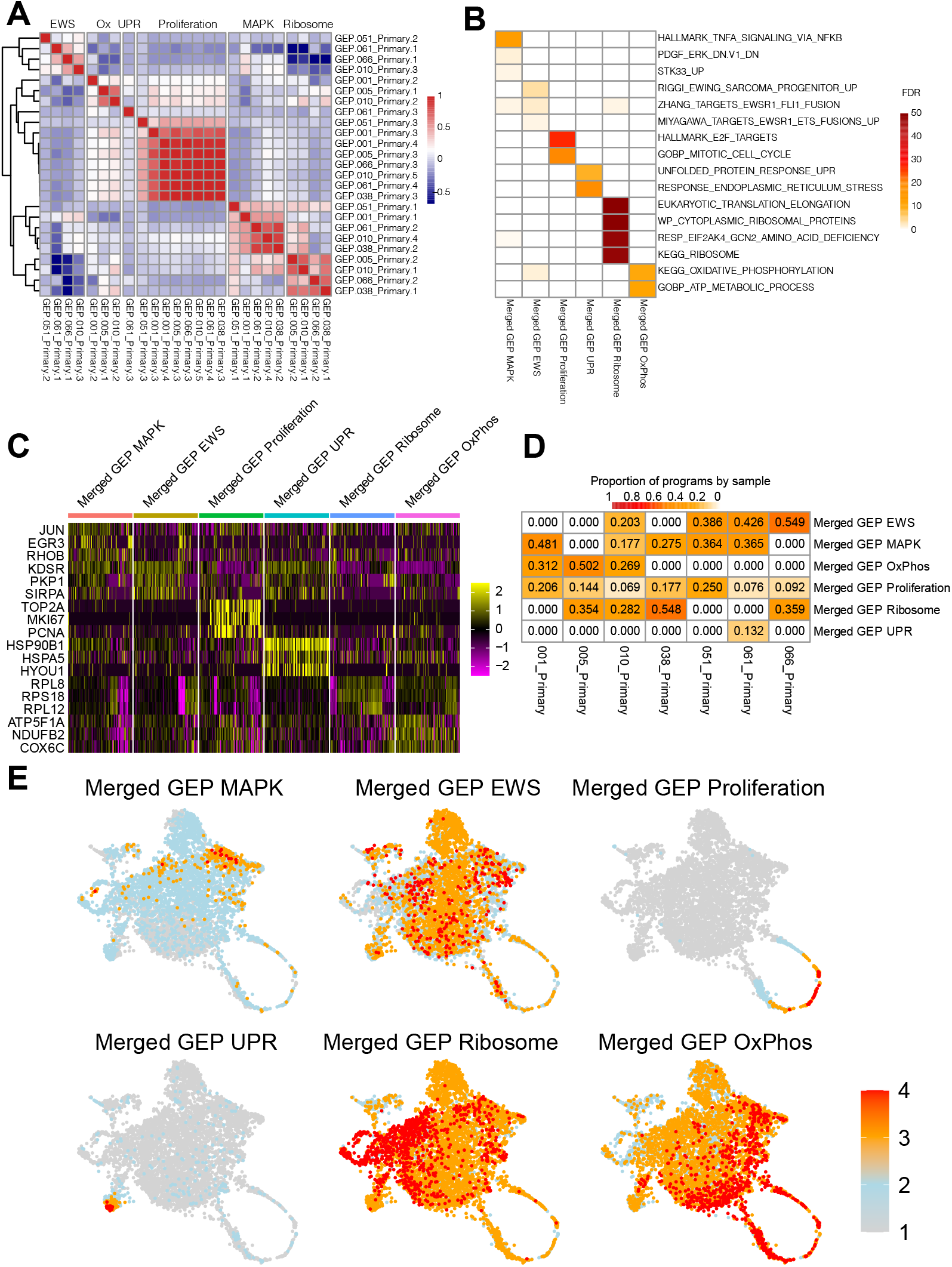
Identification of gene expression programs in Ewing sarcoma primary tumors. Correlation matrix of the full dataset scores from each sample-level gene expression program (GEP) (A). Merged GEPs analyzed using over-representation analysis and the FDR of select gene sets are displayed by program (B). Heatmap of select genes with groups downsampled to 100 cells, with yellow indicating higher expression (C). Proportion of each GEP by sample (D). Cells labeled by the GEP score (E).

Potential regulators and associations with previous gene programs were further investigated. Transcriptional drivers of each program were explored with VIPER^43^. Several drivers were as expected such as JUNB, FOXP1, E2F2, EPAS1, and ATF6 associating with merged GEP MAPK, OxPhos, Proliferation, Ribosome, and UPR, respectively (Supplementary Fig. 3A). The top drivers of the merged EWS GEP were identified as ONCECUT1, MEIS1, and POU5F1. While EWSR1 was not included in the regulon database, FLI1 and ERG were and both most closely associated with the merged Ribosome GEP. Interestingly, MEIS1 has been previously shown to cooperate with EWSR1-FLI to stimulate proliferation in Ewing sarcoma^44^, suggesting that fusion-associated drivers may account for some of the merged EWS GEP. We also compared the merged GEPs to previously identified programs in scRNA-seq from Ewing sarcoma xenografts^14^. Most merged GEPs closely correlated with programs in xenografts including between merged EWS GEP and the EwS program from Aynaud *et al.*^14^ *d*emonstrating the similarity in programs between primary tumors and model systems (Supplementary Fig. 3B).

### EWS and Proliferation merged GEPs correlate with patient outcome

Next, we investigated whether genes enriched in each merged GEP correlates with survival in previously published datasets. The two microarray data sets used in this analysis consisted of 44 Ewing sarcoma tumors which included 32 primary tumors, five recurrences, and seven metastases (GSE17679)^45^ and 85 primary Ewing sarcoma tumors which were collected from patients enrolled on COG and EuroEwing trials (GSE63157)^46^. GSE17679 was used as a discovery cohort where we found that patients with low merged GEP EWS scores or a high proliferation scores had significantly poorer survival (Fig. 3A-D). The other four merged did not correlate with overall survival (Supplementary Fig 4).

**Figure 3.**
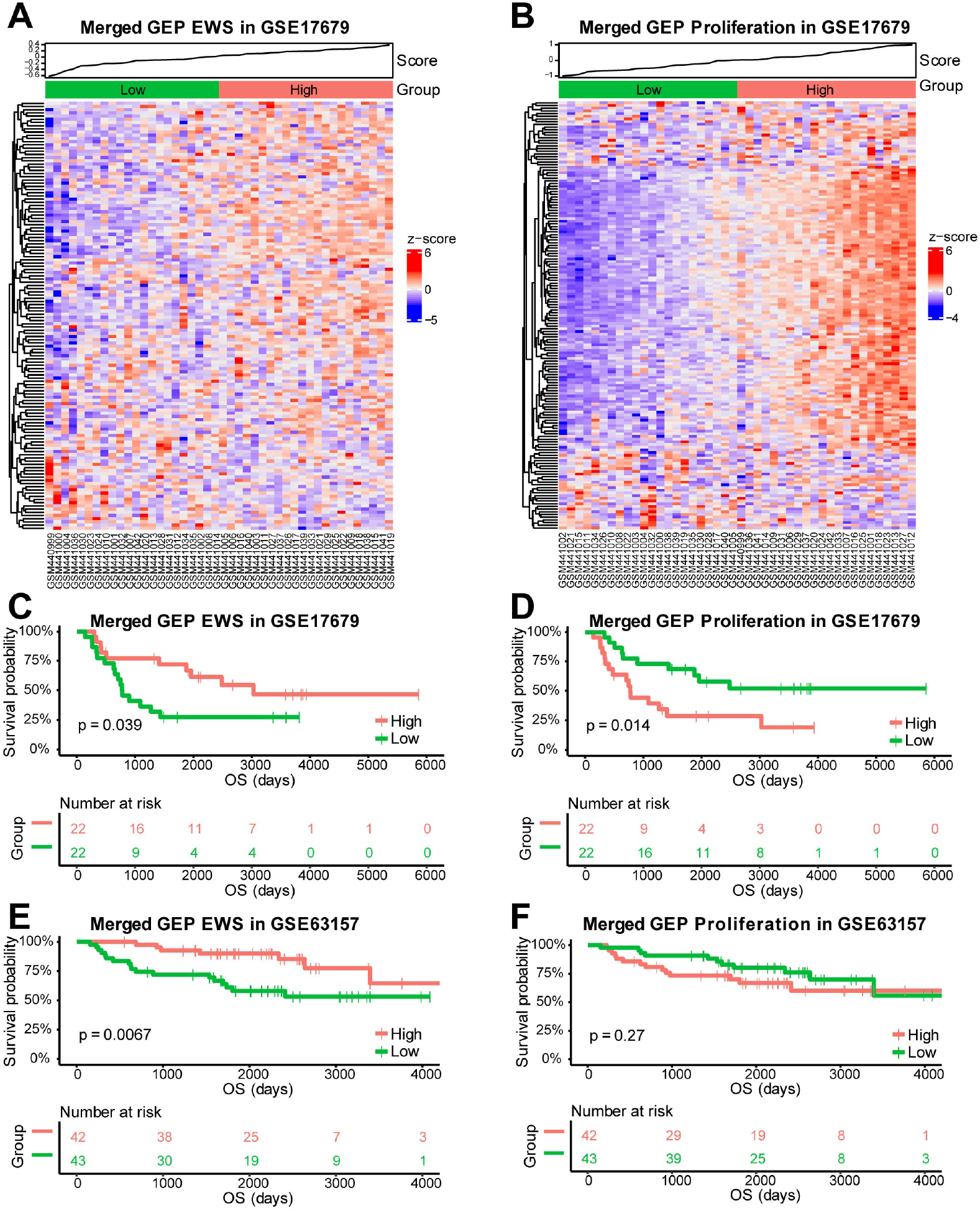
Correlation of the EWS and Proliferation gene expression programs with survival. Heatmaps displaying the expression of the genes in the EWS (A) and Proliferation (B) gene expression programs in Ewing sarcoma tumors from GSE17679. Sample scores are plotted at the top with a median split used to classify samples as low (green) or high (red) for each program. Kaplan-Meier curves with a log-rank test comparing the overall survival between the low and high groups identified from EWS (C) or Proliferation (D) gene expression programs in GSE17679 and the validation Ewing sarcoma cohort, GSE63157 (E,F).

We further evaluated merged GEP EWS and Proliferation in the validation cohort, GSE63157. Similar to GSE17679, patients with low merged GEP EWS scores had significantly poorer survival. While high merged GEP Proliferation scores correlated with poorer survival in GSE17679, it did not reach statistical significance in GSE63157 (Fig. 3E-F). There have been reports that high or low ^39^FLI1 expression in cell line models can lead to a switch between a proliferative non-motile phenotype and a motile and metastatic cell phenotype^14, 47^ but ^39^FLI1 programs have not previously associated with patient outcome. The merged GEP EWS that we have identified is a unique gene set that is enriched for EWS-FLI1 target genes which correlate to survival.

### Intratumor subclonal evolution in Ewing sarcoma

As mentioned previously, we identified that most patient tumors were found to have distinct large chromosomal gains and losses that largely agreed with clinical findings. However, other than a report utilizing molecular inversion probes, there has been little published on CNA subclones within Ewing sarcoma^48^. We leveraged scRNA-seq to analyze individual cells to identify and investigate Ewing sarcoma subclones, which were found in tumors from patients 005, 038, and 061.

In patient 005, two subclones were found to diverge following an early chr8 gain (Fig. 4A-B). One subclone, termed Subcluster.1.1, emerged with gains in chr1 and chr5 and a loss of chr11p while a smaller cluster emerged with loss of chr1p and chr16q and gains in chr2 and chr 9, suggesting both chromosomal gains and losses contribute to tumor evolution. These two subclones demonstrate significant transcriptional heterogeneity (Fig. 4C). In particular, Subcluster.1.2 expressed higher levels of CD44 and FN1 (Fig. 4D-E). CD44 is a mesenchymal stem cell marker induced upon EWS-FLI1 inhibition in A673 cells^49^ and can potentially drive Ewing sarcoma migration and invasion^50^. Interestingly, CD44 is located on chr11p13, which was not a region identified to be amplified, suggesting that subclonal evolution led to an indirect upregulation of CD44. FN1 is one of the most abundantly secreted proteins in Ewing sarcoma cell lines TC32 and CHLA10^51^, and upregulation within this subclone could be a mechanism of interaction with the tumor microenvironment by Ewing sarcoma cells.

**Figure 4.**
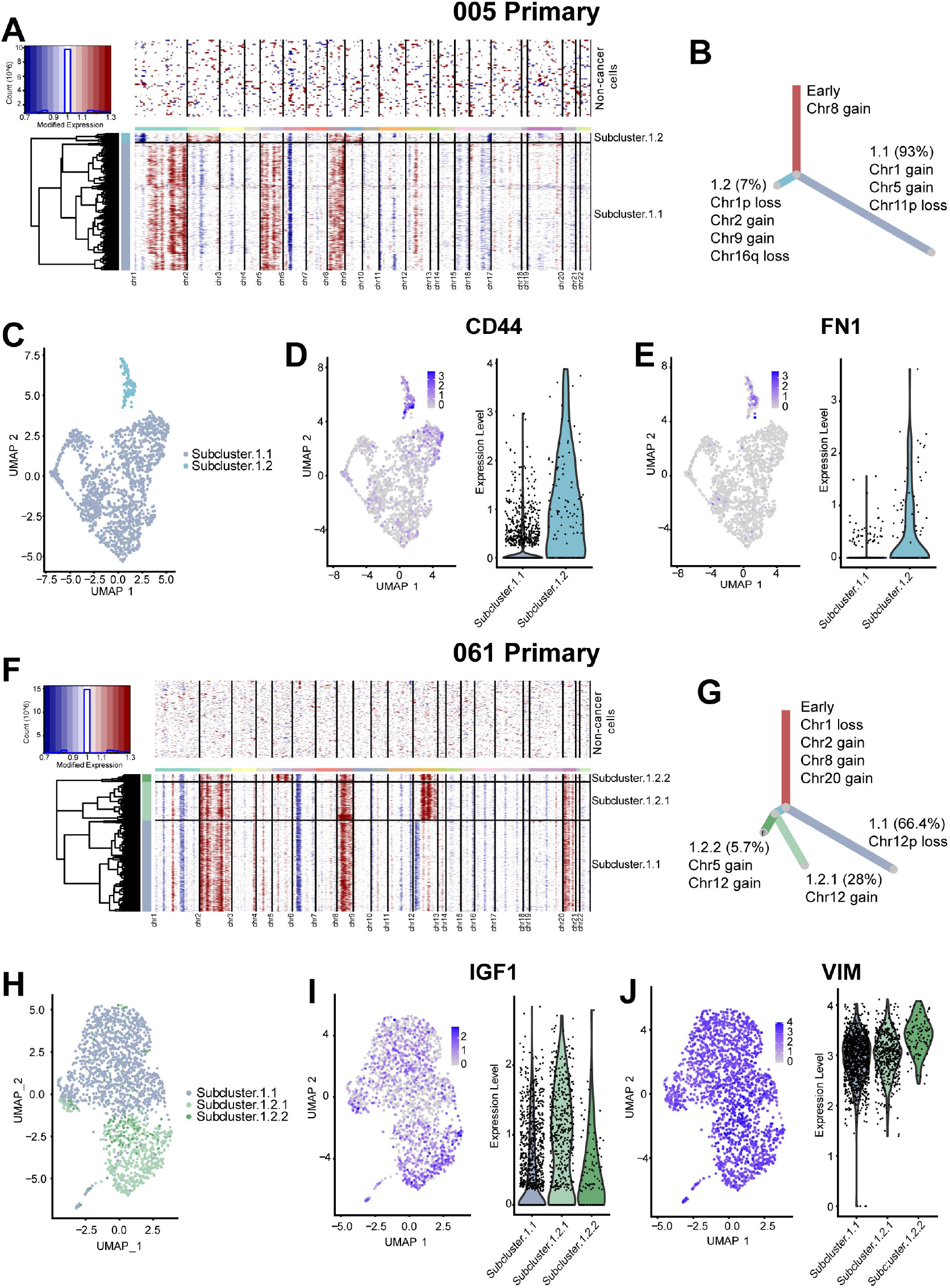
Subclones were identified in Ewing sarcoma primary tumors by analysis of copy-number alterations. Two Ewing sarcoma primary tumors displaying subclones: 005 (A-E) and 061 (F-J). Inferred copy number alterations (CNAs) in subclones of each sample (A,F). Phylogenic trees inferring evolution of the subclones based on the CNAs with the percentage of cells in each arm indicated (B,G). Subclones displayed as UMAP (C,H). Select gene expression displayed as UMAP and violin plot (D,E,I,J).

Similarly, three subclones were identified within the tumor from patient 061 (Fig. 4F). Early alterations were defined by loss of chr1 and gains of chr2, chr8, and chr20. Subclones were identified by the additional gain of chr5 and heterogenous loss and gain of chr12 (Fig. 4G). The subclones display heterogenous expression of VIM and one subclone contained higher expression of IGF1 (Fig. 4H-J). Interestingly, the diagnostic tumor chromosome analysis of the tumor from patient 061 also identified a subclone with additional gains of chr5, chr12, and chr13 (Table 1). This subclone appears to be the combination of the two minor subclones in the scRNA-seq where collectively, CNAs in chr5 and chr12 were identified. This finding validates the ability of identifying subclones in both scRNA-seq and clinical karyotyping but they may differ in resolution. The CNA analysis in the tumor from patient 038 identified three subclones but they did not lead to distinct transcriptional profiles, suggesting that not all acquired CNAs drive a differential expression profile (Supplementary Fig. 5).

### Immune infiltrates demonstrate distinct patient-dependent microenvironment phenotypes

In Ewing sarcoma, the TME has been suggested to be pro-inflammatory with high chemokine expression^52^. We investigated the composition and characteristics of the Ewing sarcoma TME, in the relatively abundant T and myeloid cells.

We were able to distinguish four subtypes within the TME, consisting of tissue resident memory, CD8+ effector, regulatory, and CD4+IL7R+ T cells (Fig. 5A, Supplementary Table 5). CD8+ effector T cells expressed CD3, CD8, KLRK1, KLRD1, and CD7^53, 54^. Notably, these cells did not express exhaustion markers such as CTLA4, TIGIT, LAG3, or PDCD1 (gene for PD1)^17^. CD4+ IL7R T cells were identified by the expression of CD3, CD4, IL7R, CD40LG, MAF, and KLRB1 (encodes CD161)^55^. The CD8+ tissue resident memory T cell cluster was differentiated by the expression of CD8, CD69, KLF6, and KLRB1^17, 56^. Finally, expression of genes such as FOXP3, BATF, and CD4 expression was used to identify regulatory T cells. The cells identified here also expressed inhibitory molecules known in regulatory T cells such as CTLA4 and TIGIT, which may be contributing to an immunosuppressive microenvironment (Fig. 5B-D)^57^. Expression patterns of several other genes also suggest an immunosuppressive environment.

**Figure 5.**
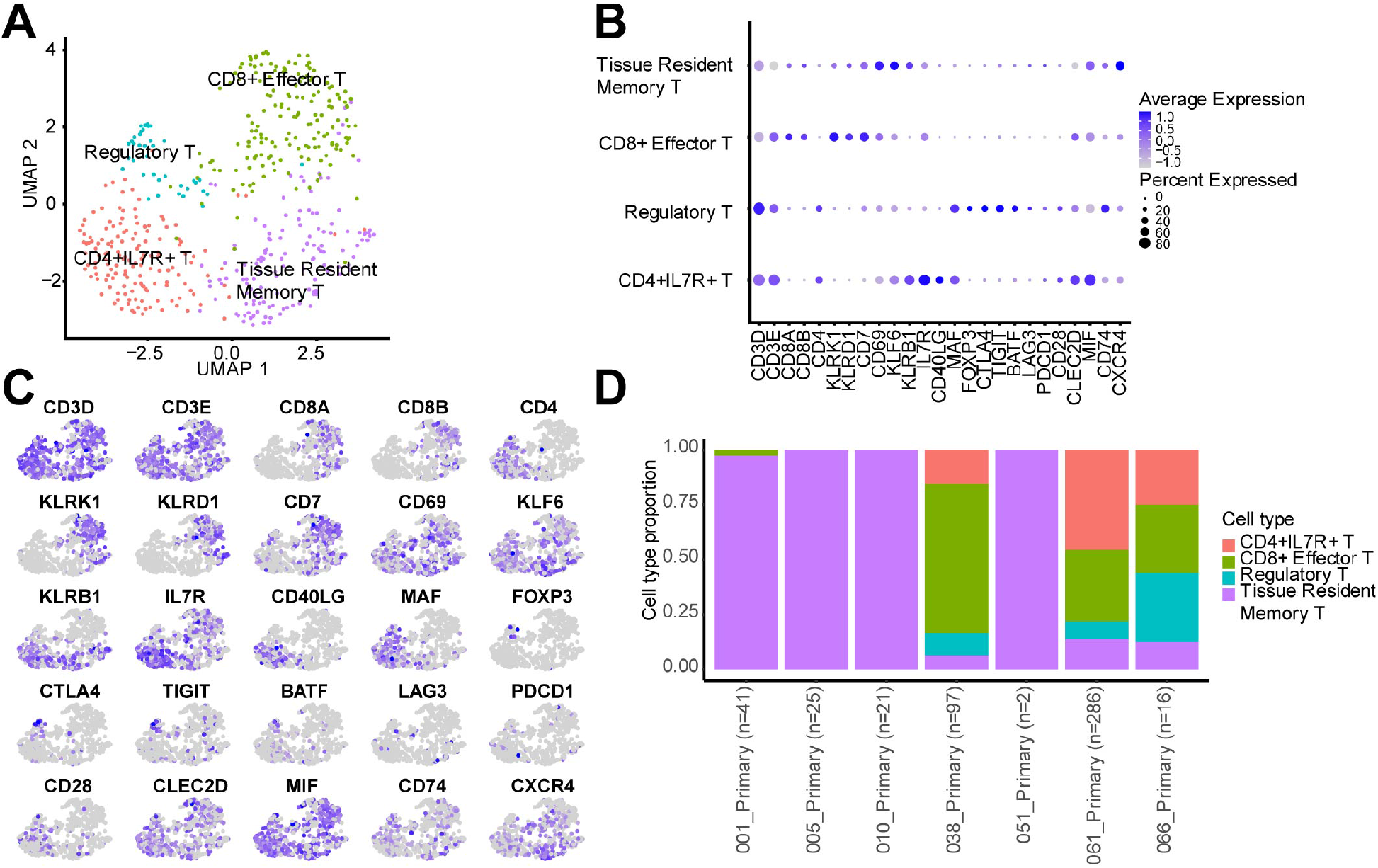
T cell subtypes in Ewing sarcoma primary tumors. UMAP showing the four identified T cell subtypes (A). Gene expression defining the subtypes displayed as a dotplot (B) or UMAP (C). Proportion of the subtypes across each sample with the number of total T cells indicated (D).

Ligands of CD161, LLT1 (encoded by CLEC2D) and CD69 (also known as CLEC2C) are expressed in the majority of the T cell subtypes in the scRNA-seq data. Most studies in immune cells suggest LLT1-CD161 interactions lead to immunosuppressive environments, although the specific relationship in memory T cells remains to be seen^58^. Additionally, CD69 has been shown to inhibit terminal differentiation of CD8+ T cells^59^. MIF expression, along with the receptors CD74 and CXCR4, are also found here in the Ewing sarcoma TME, which can lead to chemotaxis^60, 61, 62^. However, MIF binding to CD74 in T cells has been shown to reduce activation and proliferation^63^.

The myeloid population was characterized into non-inflammatory CD1C+ dendritic cells, IL1B+ inflammatory monocytes, FCN1+ inflammatory monocytes, and CD14+complement macrophages (Fig. 6A, Supplementary Table 6). The non-inflammatory CD1C+ dendritic cell cluster expressed previously reported genes for this cell type such as CD1C, CLEC10A, FCER1A, HLA-DQA1, HLA-DQB1, and HLA-DPB1^17, 64, 65, 66^. The IL1B+ inflammatory monocyte cluster is notable for the high expression of inflammation related genes such as S100A8, A100A9, S100A12, IL1B, and C5AR1, indicating this cell cluster is a pro-inflammatory monocyte population^17, 67^. The FCN1+ inflammatory monocyte cluster was identified by the higher expression of CD14, CLEC12A, CTSS, FCN1, VCAN, and LYZ, with very few cells expressing FCGR3A (gene for CD16), similar to inflammatory monocyte populations as previously reported^17, 67^. Finally, we were able to identify a CD14+ complement macrophage cluster that expresses classical macrophage related genes such as CD68, CD163, as well as complement-related genes C1QC, C1QA, and C1QB, indicating that this population is similar to previously reported C1QC+ tumor associated macrophages^17, 68^. Interestingly, these cells express APOE, PLTP, CD163, which along with C1QC indicate a strong anti-inflammatory phenotype. which could oppose the other tumor infiltrating inflammatory myeloid cells (Fig. 6B-D).

**Figure 6.**
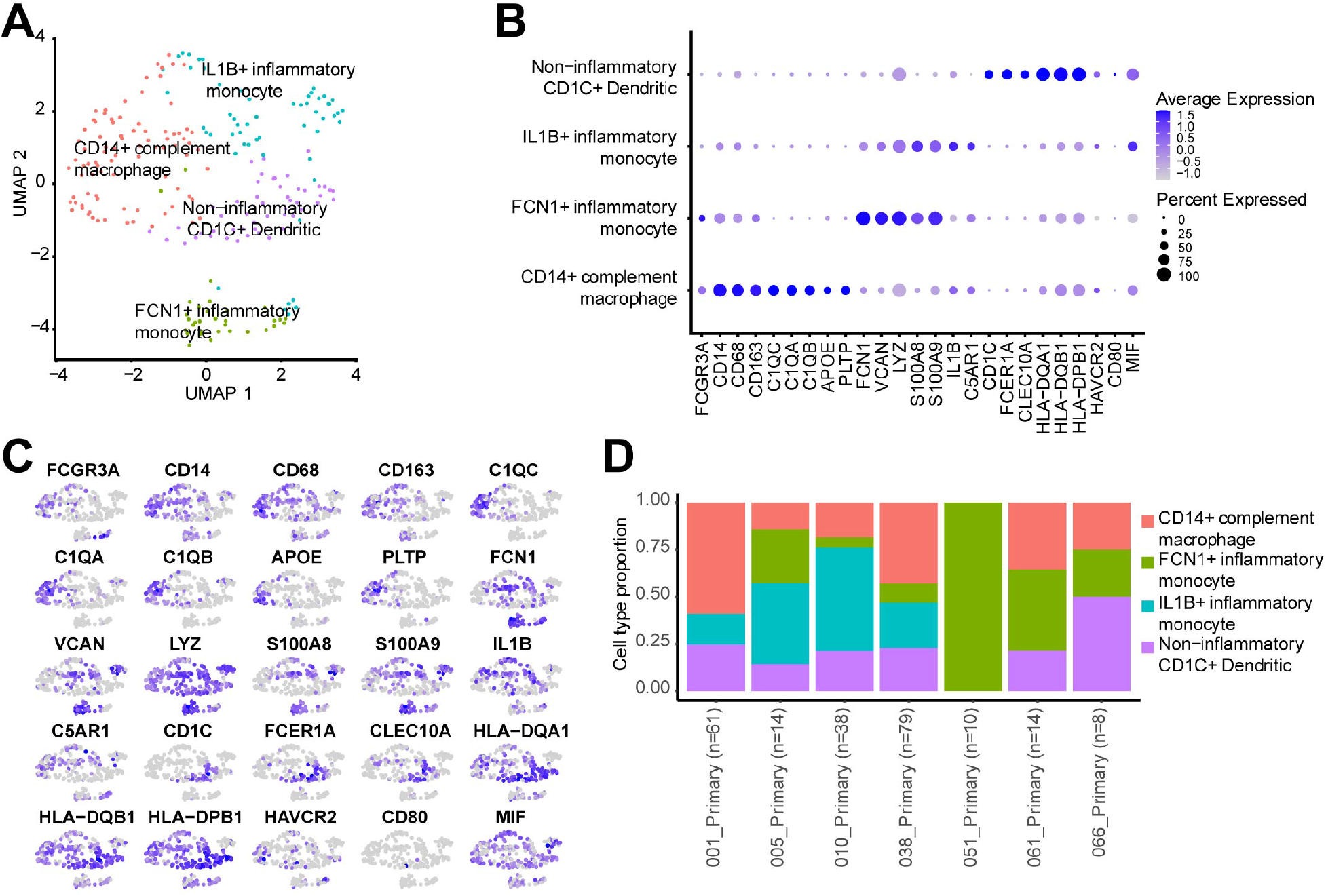
Myeloid subtypes in Ewing sarcoma primary tumors. UMAP showing the four identified myeloid subtypes (A). Gene expression defining the subtypes displayed as a dotplot (B) or UMAP (C). Proportion of the subtypes across each sample with the number of total myeloid cells indicated (D).

Similar to the T cell subtypes, the myeloid cells express several markers expected to contribute to an immunosuppressive environment. TIM3 (encoded by HAVCR2), which is moderately expressed in each of the myeloid subpopulations in this study, has been shown to reduce the response to tumors^69^. CD80, a ligand of the T cell stimulatory receptor, CD28, is lowly expressed in the myeloid cells of our study as well as those from another Ewing sarcoma study^16^ which suggests dysfunctional myeloid cells may lead to reduced T cell response. Finally, high MIF expression is similar to that in T cells which can reduce T cell activation and proliferation^63^.

### Diverse cell intrinsic and intercellular interactions are possible in the Ewing sarcoma TME

Following the determination of specific cell types in TME, we next sought to identify potential cellular interactions using CellChat^70^. Here, we were able to identify multiple ligand-receptor interactions involving Ewing sarcoma tumor cells (Fig. 7A, Supplementary Table 7). Of the tumor-intrinsic interactions, the most commonly occurring across samples were WNT5A-FZD1, NPY-NPY1R/NPY5R, and MDK-NCL (Fig. 7B). The Wnt pathway, in particular the non-canonical Wnt ligand WNT5A, has been previously reported as highly expressed in metastatic tumors, and as a potential driver of ROR1 signaling in Ewing sarcoma cells^71^. WNT5A has also been identified as a significant gene that is upregulated upon EWS-FLI1 induction in RD cells^23^. Ewing sarcoma cells have been known to highly express NPY, as well as the receptors NPY1R and NPY5R^72, 73^. MDK (midkine) has been found to be part of the Ewing sarcoma secretome^51^, however the mechanism of action, and whether NCL (Nucleolin) is a signaling receptor for midkine in Ewing sarcoma is unknown.

**Figure 7.**
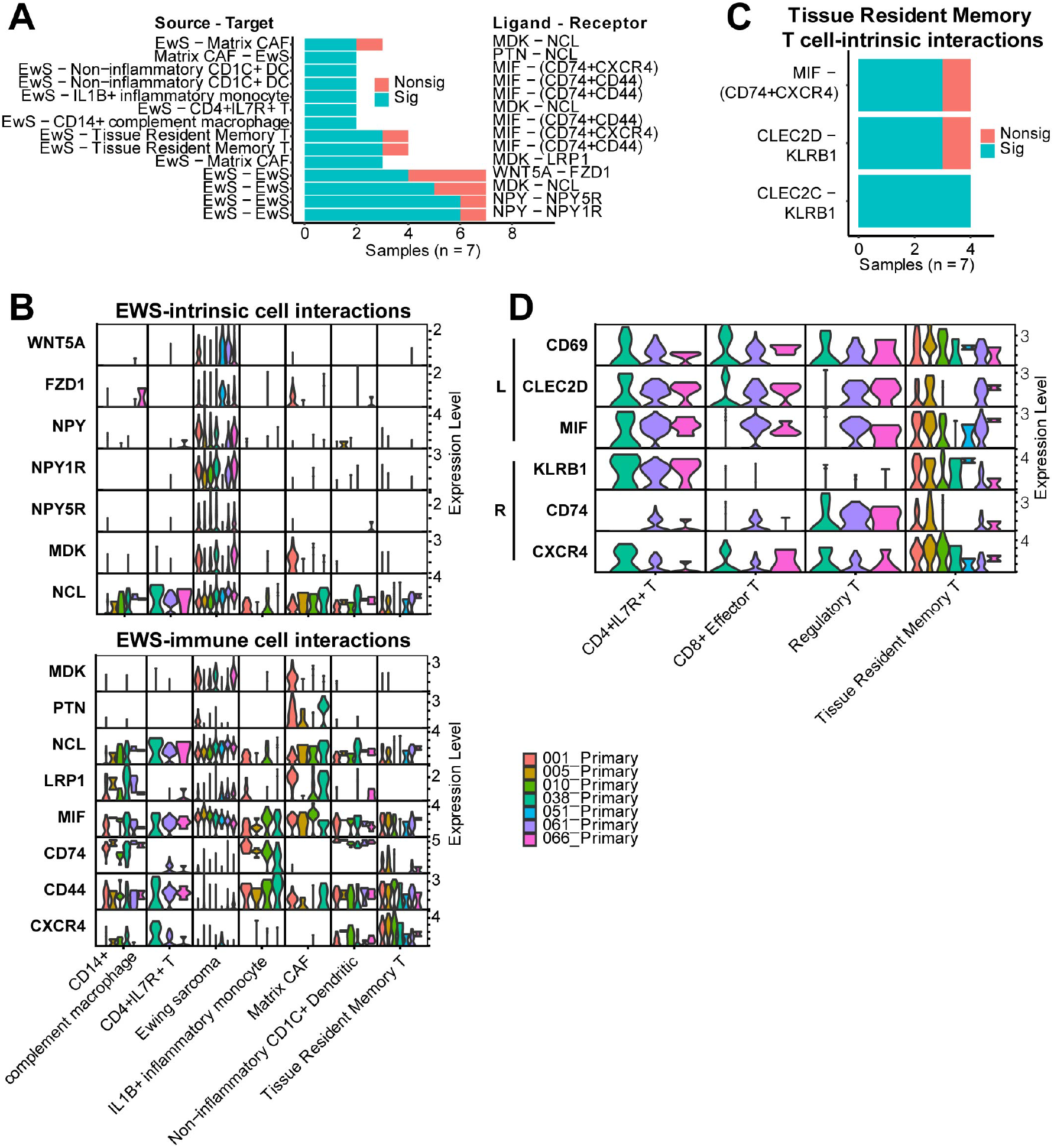
Potential cell-cell communication within Ewing sarcoma primary tumors. Select interactions involving Ewing sarcoma cells are highlighted with the number of samples where the specific interaction was significant (blue) or nonsignificant (red) (A). Not all interactions could be tested in each sample because a minimum of 10 cells were required. Expression of ligands and receptors for each sample across the cell types are plotted for Ewing sarcoma-intrinsic (top) or Ewing-immune cell interactions (bottom) (B). Significant (blue) or nonsignificant (red) interactions within immune cells in at least 3 samples (C). Ligand and receptor expression within T cell subtypes by patient sample (D).

Additionally, there are several tumor-immune interactions possible in multiple tumor samples (Fig. 7A-B). The most common across samples were interactions between MIF, expressed in Ewing sarcoma tumor cells, and CD44, CD74, and CXCR4 expressed in several different infiltrating immune cells. MIF is highly expressed across many cancer types, including glioblastoma^74^, gastric cancer^75^, neuroblastoma^76^, and osteosarcoma^77^, although this has not been reported in Ewing sarcoma. MIF was originally discovered as a cytokine that regulates inflammation and immune activation and is known to bind to the CD74/CD44 complex^78^, CXCR2 and CXCR4^79^. In mouse models of soft tissue sarcoma and melanoma, MIF has been reported to interact with CD74 expressing myeloid cells in the TME causing remodeling to a pro-tumor TME^80, 81^.

In terms of immune-intrinsic interactions, only 3 interactions were identified as probable in at least 3 patient tumor samples, all occuring within tissue resident memory T cells (Fig. 7C). MIF expression, as well as the receptors CD74 and CXCR4, are highly expressed in this cell type and as mentioned above, can promote a pro-tumor TME^80, 81^. In a cell-intrinsic manner, CD161 may be interacting with the ligands, LLT1 (encoded by CLEC2D) and CD69 (also known as CLEC2C), supporting an immunosuppressed TME^58, 59^.

### Ewing sarcoma CTCs form distinct clusters in peripheral blood samples

To further investigate mechanisms of metastatic progression, we next sought to identify and interrogate Ewing sarcoma CTCs using scRNA-seq. Eight patients had peripheral blood collected, including three patients where primary tumors were also analyzed. After collection, blood samples were enriched for CTCs using Cellsieve microfiltration membranes as previously described^82^ (Supplementary Fig. 6A). Following similar filtering to the primary tumors, 7139 cells with 9371019 UMIs across 19342 genes were identified in the blood samples. Unbiased clustering revealed ten distinct cell types, which were identified by expression of canonical cell type markers (Fig. 8A-B, Supplementary Fig. 6B, Supplementary Table 8).

**Figure 8.**
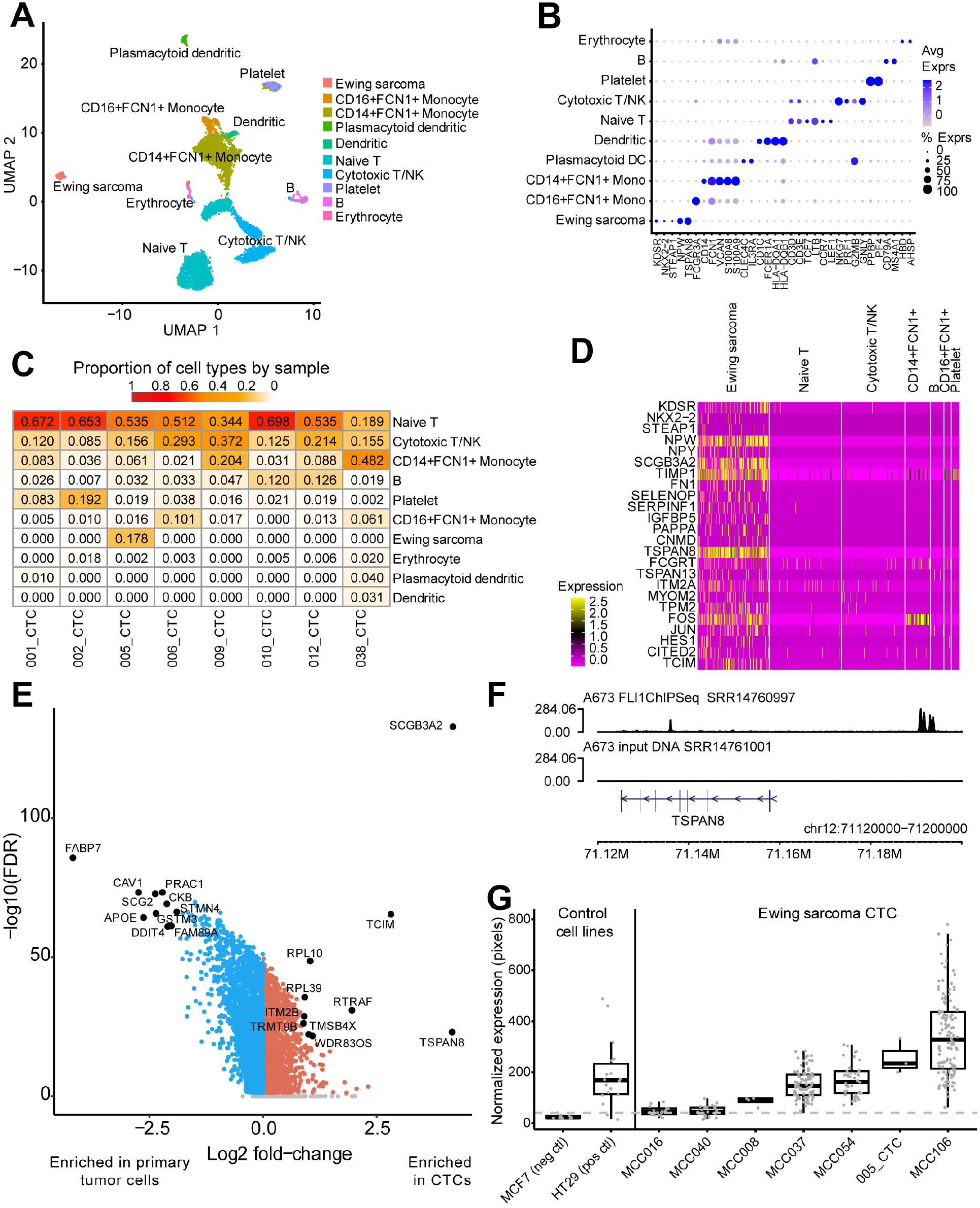
Circulating Ewing sarcoma cells identified in the blood of Patient 005. Cell types identified in the blood samples displayed as a UMAP (A) and discriminating cell markers as a dotplot (B) from all blood samples. Proportion of cell types in each sample (C). Gene expression of Ewing sarcoma-enriched genes displayed as a heatmap including only cells from 005_CTC (D). Volcano plot highlighting genes significantly enriched in 005_CTC compared to 005_Primary (E). Binding profile of FLI1 or input DNA near TSPAN8 in the A673 cell line (F). TSPAN8 staining in Ewing sarcoma CTCs. The 005_CTC sample is the same source as the 005_CTC scRNA-seq sample.

A cluster of cells were identified as Ewing sarcoma CTCs based on expression of KDSR, NKX2-2, STEAP1, and NPW. Interestingly, these Ewing sarcoma CTCs were only present in sample 005 (Fig. 8C). Since we had identified subclones within the primary tumor from this patient (Fig. 4A), we next assessed whether the CTCs represented either of those subclones. We were able to identify what appeared to be gains in chr8, chr9, and chr12 suggesting that the CTCs potentially represent both subclones found in the primary tumor (Supplementary Fig. 6C). Next, we investigated characteristic genes expressed in the identified CTCs. Recent work from the Ewing Sarcoma Cell Line Atlas (ESCLA)^83^ has identified key subsets of genes, such as core EWS-ETS binding sites, super-enhancer associated genes, and differentially expressed genes upon EWS-ETS knockdown. Here, we found that the CTCs highly expressed EWS-ETS bound genes such as FOS, CITED2, TCIM, ZNF704, PAPPA and ITM2A. Furthermore, the CTCs highly expressed CCND1, FCGRT, NRN1, TRPM4, and MYOM2, genes that were found either to be associated with super-enhancers or differentially expressed upon EWS-ETS knockdown (Fig 8D). Interestingly, one of the most highly expressed genes in the CTCs was TSPAN8, a family of the tetraspanin superfamily that has been found to be overexpressed in multiple cancers, and have been indicated to drive metastasis ^84, 85, 86, 87, 88^. TSPAN8 was also found to be highly enriched in CTCs compared to the primary tumor (Supplementary Fig. 6D-E, Fig. 8E). We examined publicly available ESCLA ChIP-seq dataand found a FLI1 binding peak proximal to TSPAN8 in all 14 cell lines (Fig. 8F, Supplementary Fig. 7), further suggesting that TSPAN8 is an EWS-FLI1 driven which may represent a therapeutic target of CTC maintenance.Importantly, Finally, to test whether TSPAN8 is expressed across otherin CTCs from other Ewing sarcoma patients, we performed TSPAN8 restaining of fixed CTCs. Diagnostic patient CTC slides containing over 20 CTCs from prospective study MCC20320 were, which expressed over 20 CTCs enumerated were selected for restaining of CTCs for restained for expression of TSPAN8 expression. Here, we were able to confirm the expression of TSPAN8 on the majority of CTCs identified across six patients, as well as from in the slide from patient 005 from our single cell RNA-seq cohort, from Ewing sarcoma stained positive for TSPAN8 protein, corroborating the scRNA-seq and suggesting TSPAN8 could be targeted on the cell surface (Fig. 8G).

## Discussion

In this work, we performed scRNA-seq on untreated Ewing sarcoma patient tumor and CTC samples to generate a dataset detailing the heterogenic copy-number and transcriptional profiles of tumor cells, as well as the composition of the TME. Previous cell line and xenograft studies have identified transcriptional programs driven by EWS-FLI1 or cell proliferation but these phenomena have not been confirmed in Ewing sarcoma patients^47, 14, 15^. With the challenges of these types of analyses in bulk RNA-seq data, our scRNA-seq dataset was interrogated for similar transcriptional programs. We also found EWS- and proliferation-driven signatures in addition to four others. Notably, GEP-EWS and GEP-Proliferation were found to correlate with survival in one microarray dataset and while GEP-EWS was significant in another, GEP-proliferation only trended towards significance. Together, these findings underscore the clinical impact of transcriptional heterogeneity.

The prognostic association of GEP-EWS is interesting because despite EWS-FLI1 and ESWR1 being well-studied in Ewing sarcoma, no related signature or protein has been previously shown to be prognostic to our knowledge. While higher activity of GEP-EWS correlating with better survival was initially surprising based on the many known pro-tumorigenic roles of EWS-FLI1, the role of this fusion is highly nuanced ^89^. EWS-FLI1 enhances metabolic and transcriptomic cell plasticity in Ewing sarcoma cells where the presence of plasticity itself may contribute to pro-tumorigenesis and treatment resistance^14, 15^. Additionally, it was recently shown EWS-FLI1 and SIX1 co-regulate an anti-metastatic gene program, which correlates with better overall survival in Ewing sarcoma^90^. Therefore, the association we observe in microarray datasets could be because of reduced metastatic potential or because lower expression of EWS-FLI1-associated genes, in a tumor where expression is typically high, may be indicative of potential for cell plasticity.

In this study, we were also able to characterize the TME of Ewing sarcoma, which contributes important aspects of tumor biology and potential therapeutic targets^91^. Priming of T cells against PAX3^92^, STEAP1^93^, and PAPPA^94^ have shown preclinical effect against Ewing sarcoma cells. PAX3 and PAPPA, but not STEAP1, were uniquely and highly expressed in tumor cells, demonstrating these could be a potential therapeutic target. While we recognize that mRNA expression does not always correlate to cell surface protein expression, we were surprised to find that many potential immunotherapeutic targets currently investigated, such as CD276 (gene for B7H3)^95^, CD99^96^, ROR1^97^, and IGF1R^97^, had significant heterogeneity within and across patient tumors at a single cell definition; this heterogeneity could play an important role in the success of related treatments.

Much of what is known about the Ewing sarcoma TME is that it is often highly immunosuppressive and this study corroborates many of those findings. We found tumor cells to express only low levels of HLA genes, consistent with previous work showing a lack of HLA expression by Ewing sarcoma cells, potentially leading to decreased T cell recognition^18^. We also found evidence that the myeloid cells of the Ewing sarcoma TME may limit T cell activation through expression of TIM3 and the low expression of CD80^69,16^. The T cell subtypes themselves also seem to support immunosuppression through LLT1-CD161, CD69-CD161, and MIF-CD74/CXCR4 interactions which all have previous evidence of supporting pro-tumor environments^58,59, 63^.

The two CAF populations we identified in our study, termed matrix and inflammatory, are similar to CAF subpopulations found in Ewing sarcoma^16^ and other cancer types including triple-negative breast^98^ and pancreatic cancer^99^. COL1A1+ CAFs have been shown to remodel the extracellular matrix which can contribute to tumor invasion and progression in breast cancer^100, 101^. Inflammatory CAFs that secrete CXCL12, which is highly expressed in the inflammatory CAFs in our study, recruit regulatory T cells, which is another mechanism to support an immunosuppressive environment^102^. However, the specific roles of these two CAF populations are not well-characterized in Ewing sarcoma and whether they perform similarly as in other cancer types remains to be seen. Taken in aggregate, our scRNA-seq analysis of the TME largely agrees with previous studies that the Ewing sarcoma TME is significantly immunosuppressed.

Historically, detecting and analyzing circulating Ewing sarcoma cells has been challenging because of low expression of markers used in typical carcinoma CTC detection, such as the epithelial cell adhesion molecule EpCAM^103, 104^. To overcome this, there have been several published approaches. First, using surface expression of CD99, flow cytometric analysis has been used to demonstrate that peripheral blood mononuclear cells from patients with Ewing sarcoma can have a higher number of CD99+CD45-mononuclear cells, compared to control^105, 106^. However, there are known populations of early monocytic progenitor cells within the bone marrow, which has made it difficult to isolate and molecularly characterize Ewing sarcoma CTCs due to high background cell contamination^107^. Ewing sarcoma cells have been found to express cell-surface vimentin, and flow cytometric isolation of Ewing sarcoma CTCs by this marker has been reported to be prognostic^108, 109^. Finally, utilizing the significant size difference of sarcoma cells to leukocytes, size-based CTC detection approaches have been performed using platforms such as ISET and CellSieve^82, 110^. We have previously reported that CellSieve can detect CTCs in many patients with high-grade sarcoma, which also includes Ewing sarcoma patients^82, 111^. In our report, we were able to molecularly characterize a population of Ewing Sarcoma CTCs for the first time. Unexpectedly, we were able to identify TSPAN8 as one of the most highly expressed genes in the CTCs. TSPAN8 has been investigated as a driver of metastasis and invasion in other cancer types^85, 87, 112, 113^, although there have been no studies performed in Ewing sarcoma. Furthermore, TSPAN8 has also been investigated as a potential cell surface target in other cancer types ^114, 115^. Future studies should explore TSPAN8 as an antibody- or CAR T cell-based therapeutic target, as well as further investigation into the role of EWS-FLI1 regulated TSPAN8 expression in tumor metastasis.

## Materials and Methods

### Tumor collection and processing

Newly diagnosed Ewing sarcoma patients were pathologically confirmed at Children’s Hospital Colorado and consented to institutional review board-approved protocol, (COMIRB 00-206) where tumor and blood samples were collected. Tumor analysis was performed under institutional review board-approved protocol, COMIRB 22-1456. Untreated primary tumor samples were biopsied and immediately preserved using DMSO:FBS(10:90) medium in a Corning CoolCell LX container chilled -1°C/minute until reaching - 80°C, followed by archival storage at -150°C. Frozen tumor tissue was thawed at 37°C and gently dissociated using an optimized collagenase-IV/DNaseI/RPMI digestion protocol. Tumor-digest suspensions were strained through a 70µm filter membrane & washed, then treated using the ErythroClear™ Red Blood Cell Depletion Kit (STEMCELL Technologies) followed by DAPI-exclusion FACS (Beckman Coulter - MoFlo™ XDP). The enriched-tumor single cell suspensions were each constructed into barcoded cDNA libraries using the Chromium Next GEM Single Cell 3’ platform (10x Genomics). Target recovery was established at 4,000 cells maximum for GEM capture and single cells were sequenced at a targeted depth of 50,000 read pairs per cell.

### Blood collection and processing

At time of diagnosis, peripheral whole blood was collected using EDTA(K2) BCTs and immediately passed through an 8µm-pore microfilter membrane (CellSieve) at a rate of 5mL/minute on a KD Scientific LEGATO 110 syringe pump. Captured cells were washed from the membrane using RPMI1640-10%FBS and immediately preserved with DMSO:FBS(10:90) medium in a Corning CoolCell LX container chilled -1°C/minute until reaching - 80°C, followed by archival storage at -150°C. Frozen cells were thawed at 37°C and treated using the ErythroClear™ Red Blood Cell Depletion Kit (STEMCELL Technologies). Cells were constructed into barcoded single-cell cDNA libraries and sequenced as described above.

### scRNA-seq processing

Cell Ranger (v6.0.1)^116^ was used to process fastq files to cell and gene count tables using unique molecule identifiers (UMIs) aligning to GRCh38 (compiled by 10X Genomics as 2020-A). Downstream quality control and analysis utilized the Seurat (v4.0.4)^117^ pipeline. Cell Ranger-filtered data was further processed by removing genes identified in fewer than 10 cells. Cell barcodes were removed if they had greater than 50,000 UMIs, greater than 7,000 genes, greater than 20% of UMIs from mitochondria, or fewer than 200 genes. Cell doublets were identified and removed using the scDblFinder R package^118^. This filtering criteria resulted in 23,547 cells across all samples totaling 22,639 genes.

The primary and CTC-enriched blood samples were further processed independently. The gene counts were normalized using Seurat by dividing each cell’s total counts and multiplying by 10,000 followed by natural-log transformation. The top 2,000 most variable genes were scaled with total UMI and percentage mitochondria regressed out. Clusters were identified as cell types through multiple methods. The top 100 enriched genes by cluster were used as input for over-representation analysis^119^ with gene sets from the MSigDb C8 collection^120^, the PanglaoDb^121^, and select Ewing sarcoma-related gene sets. Similar enrichment analysis was performed for further indicated comparisons. The SCEVAN R package was used to confirm the identification of tumor cells^122^. Canonical cell type markers were plotted in UMAP space and using dot plots. Myeloid and T cells in the primary samples were subset and analyzed separately.

### Copy-number alterations in scRNA-seq

Copy-number alterations were estimated in primary and CTC-enriched blood samples using the inferCNV R package^123^. In each data subset, cells defined as Ewing were input as the tumor while all other cells were identified as normal. Following analysis on the dataset of primary samples, three samples were analyzed further as they appeared to have subclusters. Discrete subclusters were identified using random trees whose branches were manually trimmed to fewer clusters according to the hierarchy. Phylogenetic trees were produced with the uphyloplot2 R package with major chromosomal alterations labeled^124^.

### Non-negative matrix factorization in scRNA-seq

Intratumor gene expression programs (GEPs) were identified using consensus non-negative matrix factorization (cNMF)^125^. The Ewing cells of each primary tumor were processed separately including only genes expressed in each sample (12000 genes). cNMF was run per sample with the 2000 most variant genes using raw counts first with k ranging from 2-10 and 50 iterations to select a k per sample with high stability and a low error rate. After k selection, each sample was run a final time with 200 iterations.

Sample-level GEPs were defined as the top 50 correlated genes per program. Ewing cells from all samples were scored using AddModuleScore^117^ followed by z-score transformation of each program’s score and labeling the top program per cell as the program with the highest score. Merged GEPs were identified by correlating sample-level programs using the cell scores. Each merged GEP was defined by the aggregate of genes from the appropriate sample-level programs. Genes spanning multiple merged GEPs were removed. Merged GEPs were defined by over-representation analysis as described above^119^. Cells were scored by merged GEPs as described above. UMAP plots were visualized after integrating the samples with the RPCA method^117^.

The VIPER method^43^ was run on the cell-level merged GEP labels to identify transcriptional associations from the high confidence transcription factors (A, B, and C) in the DoRothEA database^126^. Top regulators were identified by those with the highest difference in mean enrichment between a merged GEP and the rest of the cells. Cells were also scored using top independent components identified in Ewing sarcoma xenografts^14^. These scores were compared to those from merged GEPs using Pearson’s correlation.

### Evaluation of merged gene expression programs in Ewing sarcoma microarray datasets

Two publicly available Ewing sarcoma microarray datasets were analyzed using the merged GEPs. GSE17679 cohort B contains 44 samples and served as the discovery cohort while GSE63157 served as the validation cohort containing 85 samples^45, 46^. Both datasets were normalized with rma^127^. The most variable probeset was chosen for each gene. The genes for each merged GEP were used to subset the GSE17679 dataset. Z-score transformation across each gene was performed and the samples were scored by the mean expression of the transformed values. Samples were divided into two groups using the median score for each merged GEP. Kaplan-Meier plots and log-rank tests were performed using the survminer and survival R packages^128, 129^. Heatmaps were generated with the ComplexHeatmap R package^130^. The merged GEPs demonstrating a correlation with survival were tested in a similar manner in GSE63157.

### Cell communication in scRNA-seq

Cell communication between ligands and receptors in the cell types of each primary sample were predicted using the CellChat R package^70^. Select interesting interactions were highlighted based on the significant interactions across multiple samples. Ligand and receptor expression was plotted across cell type by patient sample.

### FLI1 ChIP-seq in Ewing sarcoma cell lines

Publicly available FLI1 ChIP-seq data in Ewing sarcoma cell lines (GSE176400) were analyzed for binding near the gene TSPAN8^131^. Illumina adapters and low-quality reads were filtered out using BBDuk (v.38.87) (http://jgi.doe.gov/data-and-tools/bb-tools). Bowtie2 (v.2.3.4.3)^132^ was used to align the sequencing reads to the hg38 reference human genome. Samtools (v.1.11)^133^ was used to select the mapped reads (samtools view -b - q 30) and sort the bam files. PCR duplicates were removed using Picard MarkDuplicates tool ^134^. The normalization ratio for each sample was calculated by dividing the number of uniquely mapped human reads of the sample with the lowest number of reads by the number of uniquely mapped human reads of each sample. These normalization ratios were used to randomly sub-sample reads to obtain the same number of reads for each sample using samtools view -s. Bedtools genomecov was used to create bedgraph files from the bam files^135^. Normalized Bigwig files were created using deepTools bamCoverage^136^ and visualized using the trackplot R package^137^.

### Whole-genome sequencing

Whole-genome sequencing was performed on the 7 primary tumor samples. Fastq files were processed with the nf-core sarek pipeline^138^. Mutations were evaluated using mutect2 and annotated using Ensembl VEP^139^. Genes and mutations were filtered to those previously identified in multiple Ewing sarcoma samples^5, 6^ and those that pass mutect2 and VEP filters^140^. EWSR1 translocations were identified using TIDDIT^141^ followed by annotation with SnpEff^142^. An oncoprint was generated using the ComplexHeatmap R package^130^.

### TSPAN8 staining in CTCs

Archival fixed CTC slides for TSPAN8 restaining were obtained from the National Pediatric Cancer Foundation study MCC20320. 25 patient slides were availablefor restaining, and After filtered CTCs were stained and imaged, samples were removed from storage and demounted. Filters with CTCs then underwent the QUAR-R technique as previously described^143^. Briefly, slides were soaked in PBS and filters were demounted. Filters were washed, incubated with borohydride solution, washed, and then incubated with Tris^143, 144^. After incubation, the filters were washed and stained with anti-TSPAN8 tagged with Alexa Fluor 488 at 2μg/mL (R&D Systems clone 458811). The stain was applied to the filters for 1 hour. Filters were washed and mounted on a fresh microscope slide with DAPI. CTCs were then re-read utilizing an Olympus BX54WI fluorescent microscope with Carl Zeiss AxioCam. Zen2011 Blue software was used to measure the TSPAN8 expression on each cell as previously described^143, 144^.

## Author contributions

Conceptualization: MH

Data curation: AG, ED, DLA, MH Funding acquisition: MH

Investigation: AG, AB, ABD, ED, SNT, NO, TP, RS, DW, DLA Methodology: AG, AB, ED, SNT, NO, TP, RS, DW

Project administration: MH

Resources: MH, AT, JB, ND, CC, TG, BG, SLF, JS

Software: AG, ED Supervision: MH Visualization: AG, ED

Writing—original draft: MH, AB, AG Writing—review & editing: MH, AG

## Data and materials availability

All data are available in the main text or the supplementary materials. The University of Colorado Office of Regulatory Compliance will manage and share the raw data upon request on the University share data portal. Processed data will be deposited in a separate data sharing repository upon publication. The data containing Ewing sarcoma bulk RNA seq data was downloaded from GEO (GSE17679, GSE63157).

## Supplementary Tables

Supplementary Table 1. Sequencing information of each scRNA-seq sample.

Supplementary Table 2. Significant gene enrichment by broad cell type in primary tumors.

Supplementary Table 3. Genes of each sample-level gene expression program.

Supplementary Table 4. Genes of each merged gene expression program.

Supplementary Table 5. Significant gene enrichment by T cell subtype in primary tumor.

Supplementary Table 6. Significant gene enrichment by myeloid cell subtype in primary tumors.

Supplementary Table 7. Significnat cell communication interactions involving Ewing sarcoma cells in primary tumors.

Supplementary Table 8. Significant gene enrichment by cell type in blood samples.

---

**Supplementary Figure 1.**
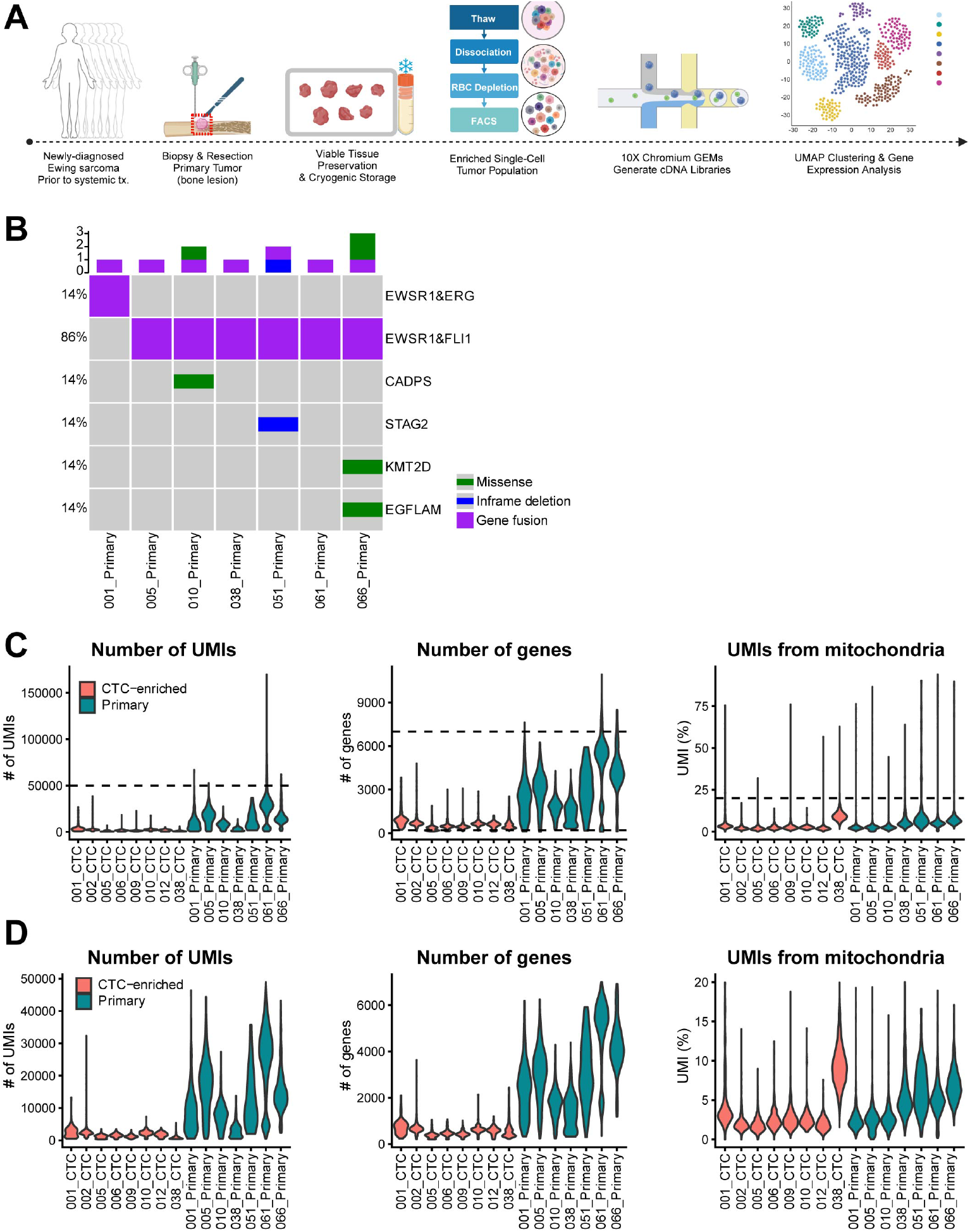
Quality control and genetic profiling of the scRNA-seq dataset. Workflow of primary tumor sample collection and processing for scRNA-seq (A). Oncoprint displaying alterations found in the primary tumors using whole-genome sequencing (B). Quality control of the scRNA-seq dataset showing the distribution of UMIs, genes, and mitochondria in barcodes considered to be cells from each sample pre- (C) or post-filtering (D). The dashed lines in A indicate the chosen filtering criteria.

**Supplementary Figure 2.**
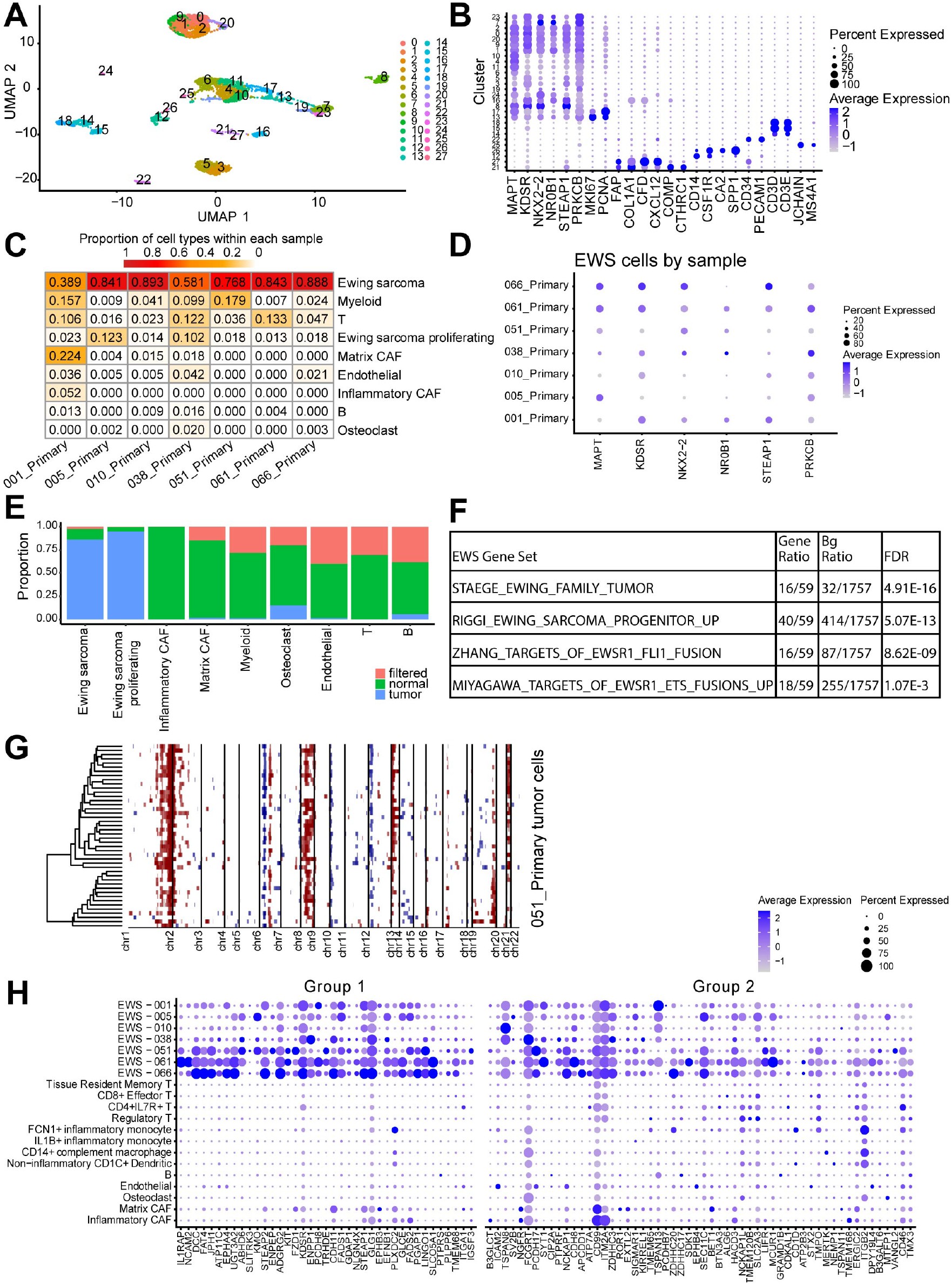
Distribution of clusters and cell types in primary Ewing sarcoma tumors. Expression of canonical marker genes across clusters are shown as a UMAP (A) or dotplot (B). Proportion of all cell types by sample (C). Expression of six canonical Ewing sarcoma-related genes in the tumor cells identified from each sample (D). SCEVAN tumor classification results distributed by the identified cell type (E). Over-representation analysis of Ewing sarcoma-related gene sets in the tumor cell clusters (F). Inferred copy-number alteration estimated in 051 tumor cells. Same as Figure 1E (G). Expression of high priority group 1 and group 2 immunotherapeutic target candidate genes by Mooney *et al*^39^.

**Supplementary Figure 3.**
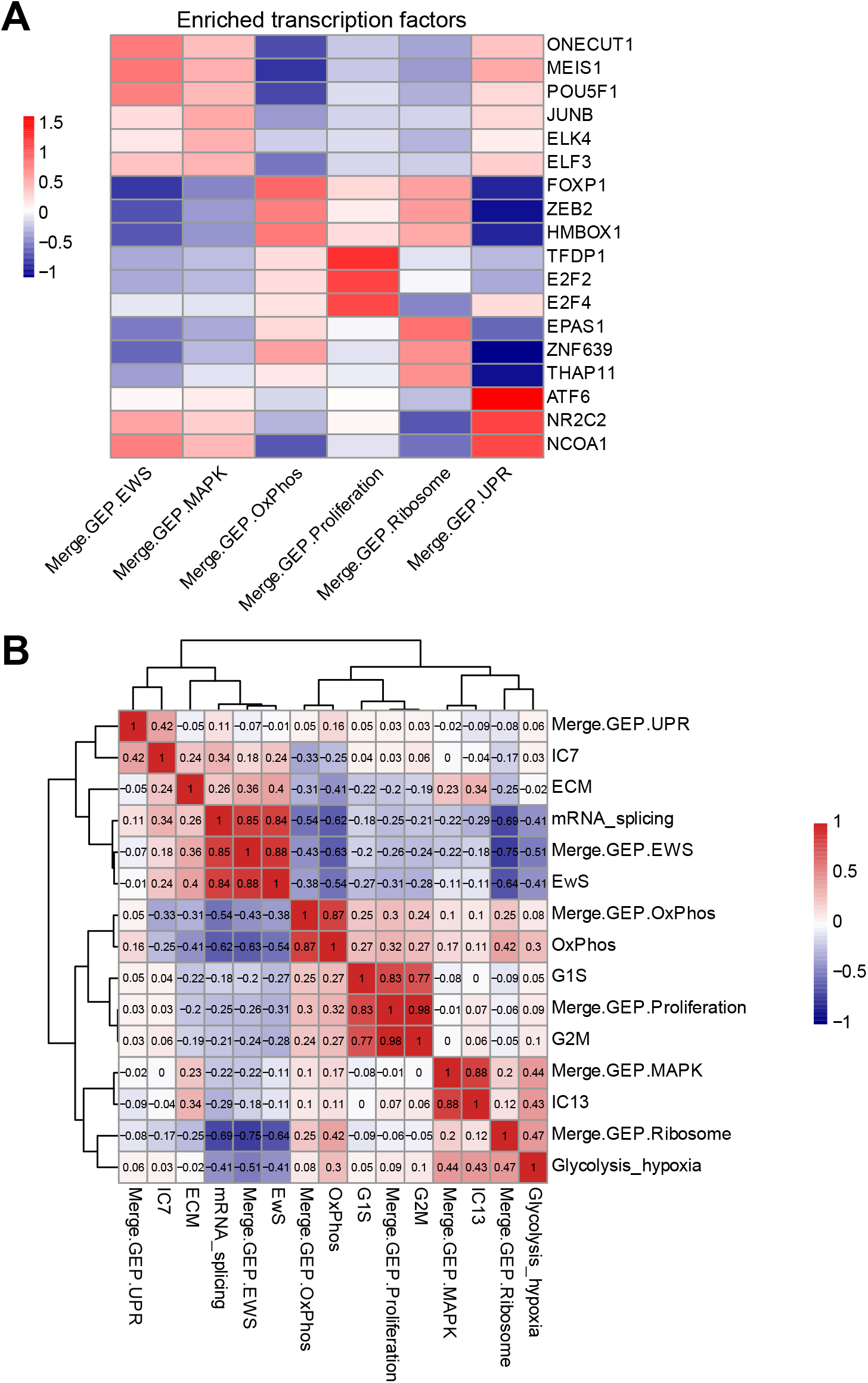
Characterization of merged gene expression programs (GEPs). Heatmap displaying the mean scaled normalized enrichment scores for the top three enriched transcriptional regulators by merged GEP (A). Correlation of cell scores between merged GEPs and programs from Aynaud *et al* (B)^14^.

**Supplementary Figure 4.**
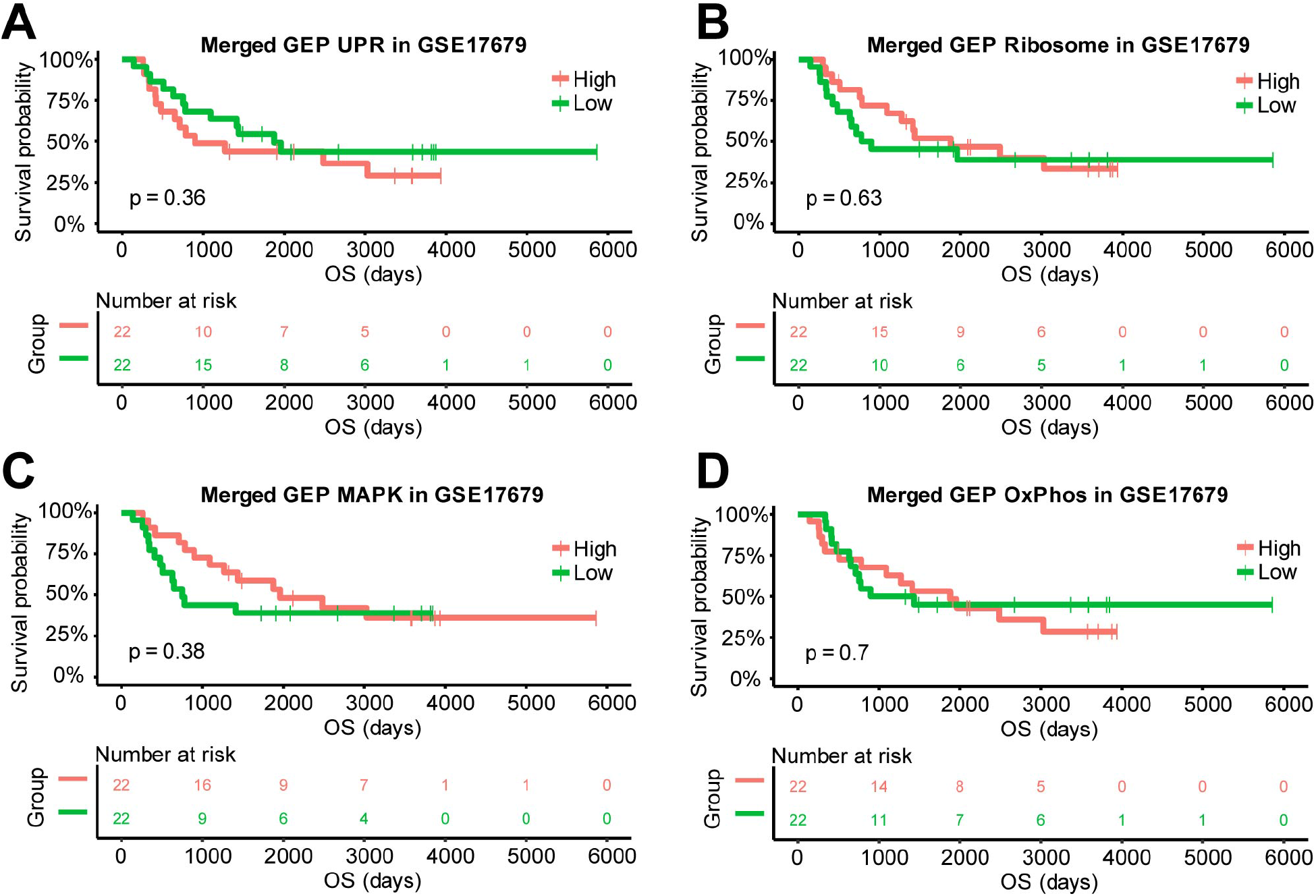
Correlation of four gene expression programs with survival in the discovery cohort. Kaplan-Meier curves with a log-rank test comparing the overall survival between the low and high groups identified from the UPR (A), Ribosome (B), MAPK (C), or OxPhos (D) gene expression programs in GSE17679.

**Supplementary Figure 5.**
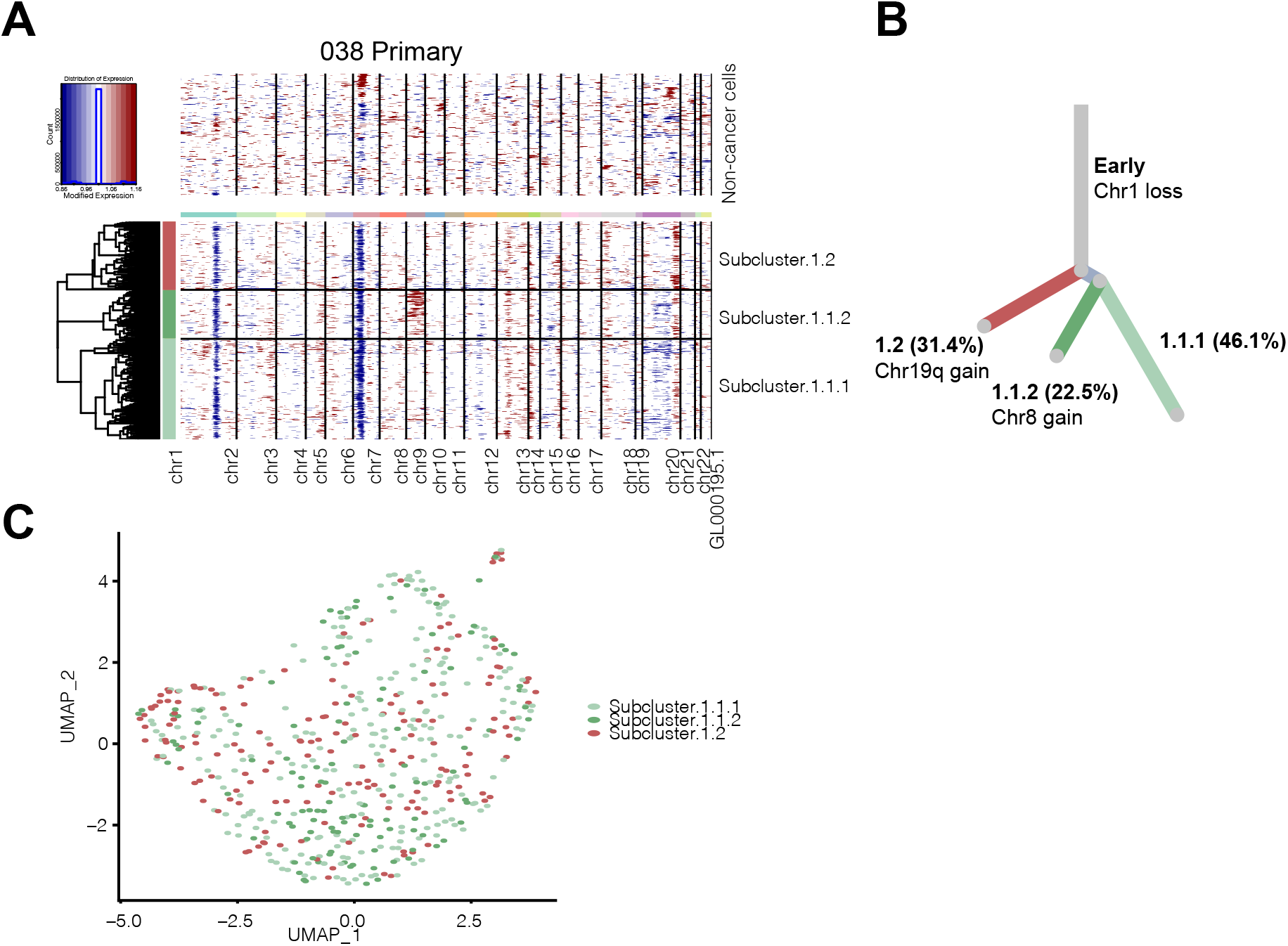
Subclones were identified in Ewing sarcoma primary tumor sample 038 by analysis of copy-number alterations. Inferred copy number alterations (CNAs) in subclones of sample 038 (A). Phylogenic tree inferring evolution of the subclones based on the CNAs with the percentage of cells in each arm indicated (B). Subclones displayed as UMAP (C).

**Supplementary Figure 6.**
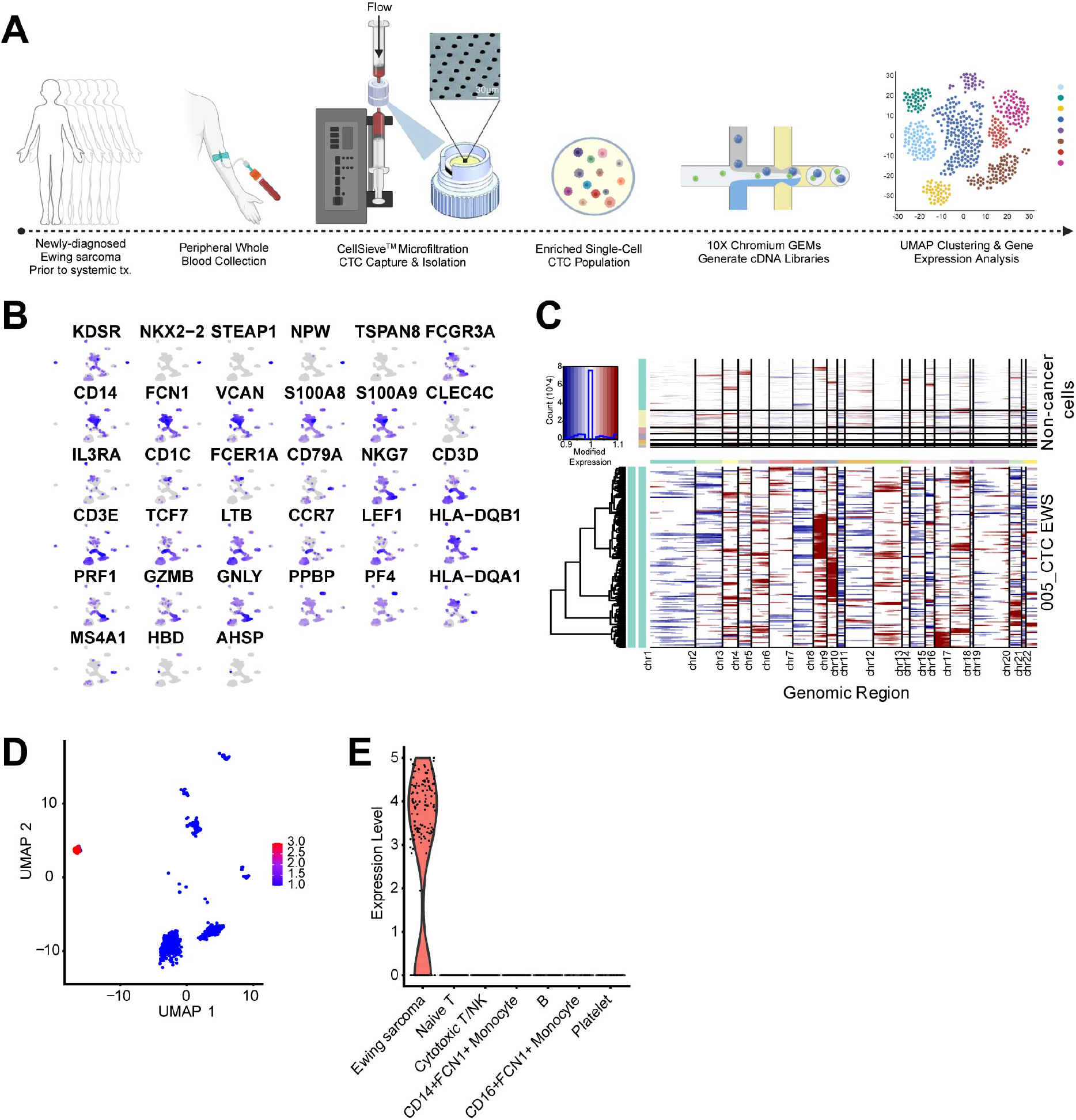
Inferred copy-number alterations and TSPAN8 expression in blood. Workflow of blood sample collection, enrichment of circulating tumor cells, and processing for scRNA-seq (A). Cell type markers (B). Inferred copy-number alterations of sample 005 Ewing sarcoma cells compared to normal cells (C). TSPAN8 expression in sample 005 as UMAP (D) and violin plots by cell type (E).

**Supplementary Figure 7.**
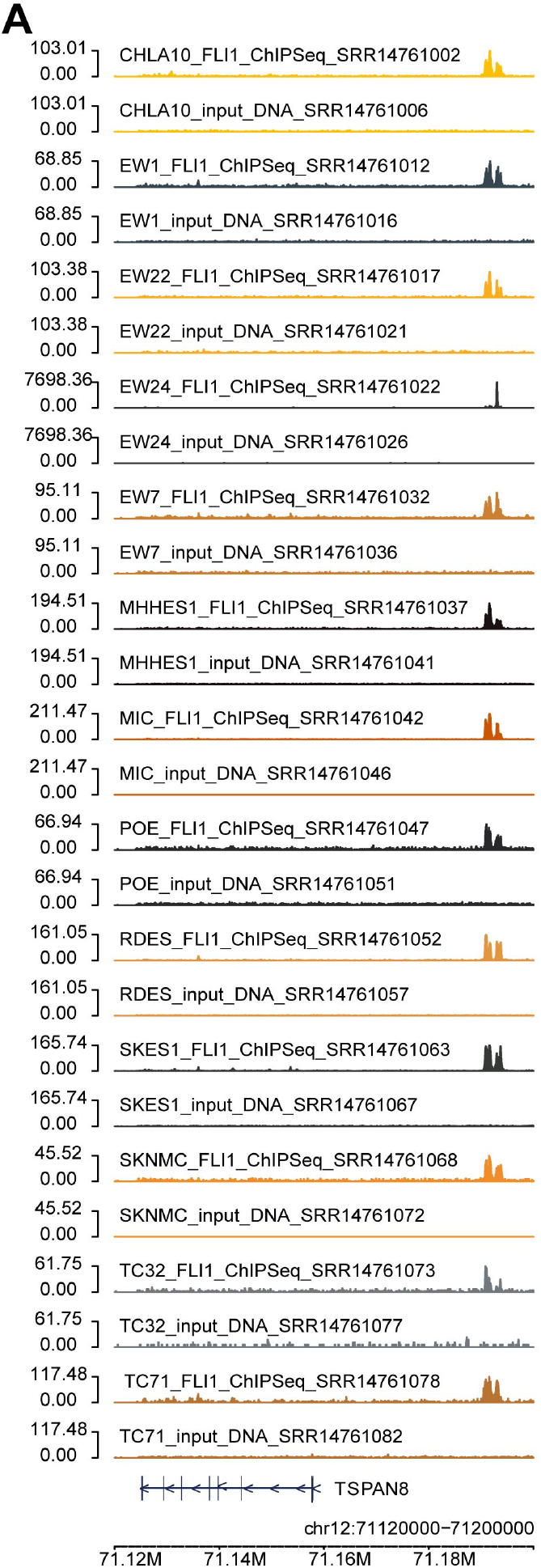
FLI1 ChIP-seq binding near TSPAN8. Binding profile of FLI1 or input DNA near TSPAN8 in several Ewing sarcoma cell lines (A).

